# Learning to select actions shapes recurrent dynamics in the corticostriatal system

**DOI:** 10.1101/646141

**Authors:** Christian D. Márton, Simon R. Schultz, Bruno B. Averbeck

## Abstract

Learning to select appropriate actions based on their values is fundamental to adaptive behavior. This form of learning is supported by fronto-striatal systems. The dorsal-lateral prefrontal cortex (dlPFC) and the dorsal striatum (dSTR), which are strongly interconnected, are key nodes in this circuitry. Substantial experimental evidence, including neurophysiological recordings, have shown that neurons in these structures represent key aspects of learning. The computational mechanisms that shape the neurophysiological responses, however, are not clear. To examine this, we developed a recurrent neural network (RNN) model of the dlPFC-dSTR circuit and trained it on an oculomotor sequence learning task. We compared the activity generated by the model to activity recorded from monkey dlPFC and dSTR in the same task. This network consisted of a striatal component which encoded action values, and a prefrontal component which selected appropriate actions. After training, this system was able to autonomously represent and update action values and select actions, thus being able to closely approximate the representational structure in corticostriatal recordings. We found that learning to select the correct actions drove action-sequence representations further apart in activity space, both in the model and in the neural data. The model revealed that learning proceeds by increasing the distance between sequence-specific representations. This makes it more likely that the model will select the appropriate action sequence as learning develops. Our model thus supports the hypothesis that learning in networks drives the neural representations of actions further apart, increasing the probability that the network generates correct actions as learning proceeds. Altogether, this study advances our understanding of how neural circuit dynamics are involved in neural computation, showing how dynamics in the corticostriatal system support task learning.

## 1 Introduction

Human and nonhuman primates are capable of complex adaptive behavior. Adaptive behavior requires predicting the values of choices, executing actions on the basis of those predictions, and updating predictions following the rewarding or punishing outcomes of choices. Reinforcement learning (RL) is a formal, algorithmic framework useful for characterizing these behavioral processes. Experimental work suggests that RL maps onto fronto-striatal systems, dopaminergic interactions with those systems, and other structures including the amygdala and thalamus [Averbeck and Costa, 2017]. Little is known, however, about how the RL formalism and the associated behaviors map onto mechanisms at the neural population level across these systems. How do neural population codes evolve with learning across these systems, and what are the underlying network mechanisms that give rise to these population codes?

Experimental work and modeling has implicated fronto-striatal systems in aspects of RL [Niv, 2009, Lee et al., 2012, Botvinick, 2012, Botvinick and Weinstein, 2014, Wang et al., 2018, Langdon et al., 2018, Niv, 2019]. Several studies have suggested that the striatum codes action values [Houk, 1995, Suri and Schultz, 1998, Doya, 1910, Nakahara et al., 2001, O’Doherty et al., 2004, Frank et al., 2004, Frank, 2005, Samejima et al., 2005, Pasupathy and Miller, 2005, Histed et al., 2009, Amemori et al., 2011, Daw et al., 2011, Sarvestani et al., 2011, Li and Daw, 2011, Seo et al., 2012, Averbeck and Costa, 2017]. These studies have further suggested that the phasic activity of dopamine, which codes reward prediction errors, drives updates of the striatal action value representations following reward feedback. Several areas in the PFC have also been implicated in action selection and decision making [Wood and Grafman, 2003, Friston, 2005, Averbeck et al., 2006, Cheong et al., 2006, Koechlin and Summerfield, 2007, Tsujimoto et al., 2008, Sakai, 2008, Histed et al., 2009, Collins and Koechlin, 2012, Botvinick, 2012, Stokes et al., 2013, Lim and Goldman, 2013, Verschure et al., 2014, Botvinick and Weinstein, 2014, Domenech and Koechlin, 2015, Rich and Wallis, 2016, Balaguer et al., 2016, Alexander et al., 2018, Aitchison and Lengyel, 2017, Wallis et al., 2019, Radulescu et al., 2019]. These studies further suggest that PFC plans future actions and predicts future outcomes. While both striatum and PFC have been found to represent action value and choice signals, value signals were found to be stronger in dSTR than lPFC, while action related signals were stronger in lPFC [Pasupathy and Miller, 2005, Samejima et al., 2005, Averbeck et al., 2006, Seo et al., 2012].

Previous work in the motor system and in prefrontal cortex has shown that insight into the computational mechanisms that underlie complex tasks can be gained by treating neural populations as a dynamical system and studying how their trajectories evolve with time [Rabinovich et al., 2008, Buonomano and Maass, 2009, Sussillo and Abbott, 2009, Sutskever, 2013, Shenoy et al., 2013, Mante et al., 2013, Sussillo and Barak, 2013, Barak et al., 2013, Hennequin et al., 2014, Carnevale et al., 2015, Rajan et al., 2016, Gallego et al., 2017, Wang et al., 2017, Chaisangmongkon et al., 2017, Remington et al., 2018, Wang et al., 2018, Yang et al., 2018, Gallego et al., 2018, Sussillo et al., 2015, Botvinick et al., 2019, Musall et al., 2019, Richards et al., 2019]. In prefrontal cortex this work has helped shed light on how task execution is driven by dynamics around fixed and slow points in neural population space [Mante et al., 2013, Chaisangmongkon et al., 2017]. These studies have examined representations and computational mechanisms in decision making tasks, where the values of choices have already been learned previously. In the present study, we have used a similar approach to study how representations *develop* as animals learn to make choices that deliver rewards.

In accordance with these findings, we built a joint recurrent network model of the corticostriatal system in which the striatal network represents RL-derived action values and the prefrontal cortex, via recurrent basal ganglia loops, selects appropriate actions based on this signal. We trained this system on a complex decision making task. We also obtained neural recordings from these two regions in two macaques trained on the same task.

We hypothesized that learning would drive specific structure in state space dynamics. We further hypothesized that a system designed to learn RL-derived action values and select appropriate actions based on them would be computationally similar to the fronto-striatal system in the brain. If so, the representational structure during learning in the network should be similar to that found in neural recordings. Moreover, the differing roles assigned to the striatal and prefrontal networks should suffice to induce a difference in representational structure across the two regions in a way that matches the asymmetries in action value and choice representation observed previously [Seo et al., 2012].

Investigating the change in representational structure with learning, we found that dynamic movement-sequence representations moved apart from each other in latent space with learning, in both the model and the neural data. We found that this process was driven by the evolution of potential surfaces in the networks such that movement-sequence specific gradient minima moved farther apart in activity space. This increase in distance, or in other words, the increase in the height of the gradient hill between sequence representations, makes it less likely that the wrong action is selected as learning proceeds.

## 2 Methods and Materials

### 2.1 Neural Data

The neural data employed here has been previously published in [Seo et al., 2012], though not with the analysis that has been carried out here.

#### 2.1.1 Subjects

Two adult male rhesus monkeys (Macaca mulatta) weighing 5.5–10 kg were used for recordings. All procedures and animal care were conducted in accordance with the Institute of Laboratory Animal Resources Guide for the Care and Use of Laboratory Animals. Experimental procedures for the first animal were in accordance with the United Kingdom Animals (Scientific Procedures) Act of 1986. Procedures for the second animal were in accordance with the National Institutes of Health Guide for the Care and Use of Laboratory Animals and were approved by the Animal Care and Use Committee of the National Institute of Mental Health (NIMH).

#### 2.1.2 Task and Stimuli

The two animals performed an oculomotor sequential decision-making task (Fig. 1A). A particular trial began when the animals acquired fixation on a green circle (Fixate). If the animal maintained fixation for 500 ms, the green target was replaced by a dynamic pixelating stimulus with a varied proportion of red and blue pixels and the target stimuli were presented (Stim On). The fixation circle stimulus was generated by randomly choosing the color of each pixel in the stimulus (n = 518 pixels) to be blue (or red) with a probability *q*. The color of a subset (10%) of the pixels was updated on each video refresh (60 Hz). Whenever a pixel was updated its color was always selected with the same probability *q*. The set of pixels that was updated was selected randomly on each refresh. In the original experiment we focused on differences between choices driven by reinforcement learning and choices driven by immediately available information, in alternating blocks of trials. In the present manuscript we will only consider the learning blocks. The pixelating stimulus was relevant for the blocks in which choices were driven by immediately available information. Therefore, we will not consider this stimulus further.

**Fig 1.**
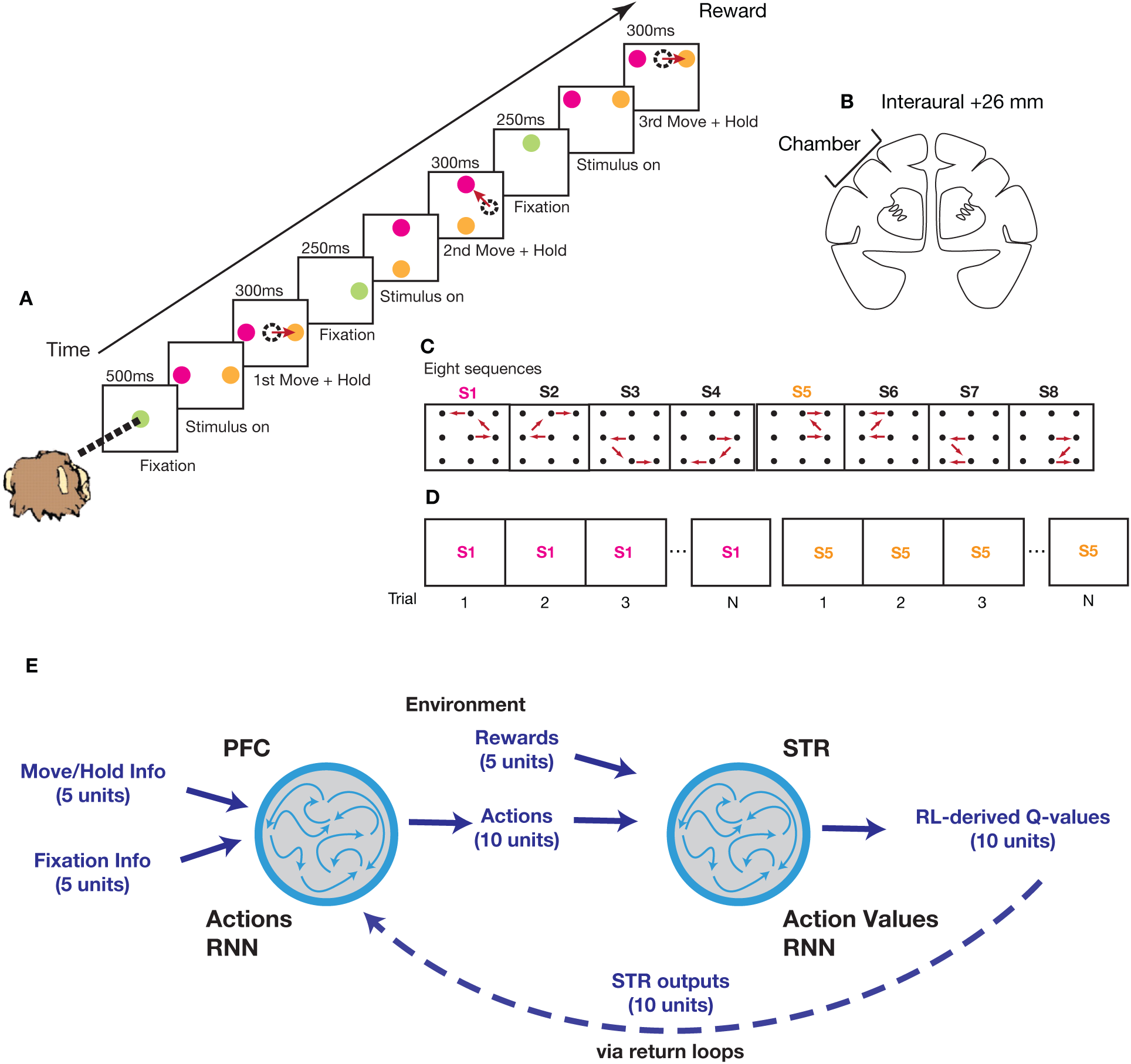
Task and Model Overview. (A) Sequence of events in the task. (B) Coronal section showing approximate location of recording chamber. (C) The eight possible movement-sequences used in the task. (D) Trial structure. Trials were arranged into blocks of 8 repeats of the correct movement-sequence plus a variable number of error trials (i.e. sequence S1 followed by sequence S5). (E) Corticostriatal model, consisting of a prefrontal network and a striatal network of recurrently connected units. The prefrontal network selects actions based on inputs from the striatal network. The striatal network outputs action values based on inputs specifying actions performed and rewards received.

The animal’s task was to saccade to the correct target (Fig. 1A). The animal could make their decision at any time after the target stimuli appeared. After the animal made a saccade to the peripheral target, it had to maintain fixation for 300 ms to signal its decision (first Move + Hold). If the saccade was to the correct target, the target then turned green and the animal had to maintain fixation for an additional 250 ms (Fixate). After this fixation period, the green target was again replaced by a fixation stimulus and two new peripheral targets were presented (Stim On). In the case that the animal made a saccade to the wrong target, the target was extinguished and the animal was forced back to repeat the previous decision step. This was repeated until the animal made the correct choice. For every trial the animal’s task was to correctly execute a sequence of three correct decisions for which the animal received either a juice reward (0.1 ml) or a food pellet reward (TestDiet 5TUL 45 mg). After that, a 2000ms inter-trial interval began. The animals always received a reward if they reached the end of the sequence of three correct decisions, even if errors were made along the way. If the animal made a mistake, it only had to repeat the previous decision, it was not forced back to the beginning of the sequence. The full task included both fixed and random conditions, as explained in detail in [Seo et al., 2012]. In the present study, however, only data from the fixed condition was used.

There were eight possible sequences in this task as every trial was composed of three binary decisions (Fig. 1C). The eight sequences were composed of ten different possible individual movements. Every movement occurred in at least two sequences. We also used several levels of color bias, q as defined above. On most recording days in the fixed sets we used *q* ∈ (0.50, 0.55, 0.60, 0.65). The color bias was selected randomly for each movement and was not held constant within a trial. Choices on the 50% color bias condition were rewarded randomly. The sequences were highly overlearned. One animal had 103 total days of training, and the other 92 days before chambers were implanted. The first 5–10 days of this training were devoted to basic fixation and saccade training.

In the fixed condition employed here, the correct spatial sequence of eye movements remained fixed for blocks of eight correct trials (Fig. 1D). After eight trials were executed without any mistakes, the sequence switched pseudorandomly to a new one. Thus, the animal could draw on its memory to execute a particular sequence, except following a sequence switch.

#### 2.1.3 Neural Data Analysis

The neural data analyzed comprised 365 units from dSTR and 479 units from lPFC (Fig. 1B, and as explained in more detail in [Seo et al., 2012]). Data from individual movements was not analyzed if animal failed to maintain fixation or did not saccade to one of the choice targets. We fitted an ANOVA model with a 200ms sliding window applied in 25ms steps aligned to movement onset, as done previously [Seo et al., 2012], and identified units that did not show a significant effect for the *sequence* factor at any point across the entire recording session. These units were excluded from the analysis, as were units that showed average firing rates below 1*Hz*. For subsequent analysis, data was pooled across animals and recording sessions and averaged across runs.

To analyze neural population responses, we applied demixed principal component analysis (dPCA) [Brendel et al., 2011, Kobak et al., 2016] to the firing rate traces. As a dimensionality reduction technique, dPCA strives to find an encoding (or latent representation) which captures most of the variance in the data but also expresses the dependence of the representation on different task parameters such as stimuli or decisions. More specifically, it decomposes neural activity into different task parameters, in our case time (**X**_*t*_), sequence (**X**_*s*_), and certainty (**X**_*t*_), and any combination of those (**X**_*sc*_, **X**_*st*_, **X**_*ct*_):

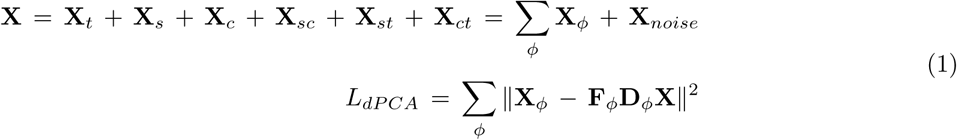

In order to obtain task trajectories in the reduced dPCA space, the data (**X**) was smoothed with a Gaussian kernel and then projected into 3-dimensional latent space spanned by the first three vectors of the *sequence*-decoder matrix (**D**_*s*_).

Distance measures were obtained on the full datasets in full-dimensional neural space, not in the reduced subspace. Euclidean distance between all sequences was computed across all time points for each of the 8 trial repeats and averaged across all possible sequence combinations. For PFC, distances were computed between sequences *within* the two clusters (defined by whether a sequence ended in the upper or lower visual hemisphere (sequences S1, S2, S5, S6 and S3, S4, S7, S8, respectively; see Fig.1C). As a measure of how compact the trajectories were, the Euclidian distance of every sequence to its centroid was computed across all time points for each of the 8 trial repeats and averaged across sequences. A sequence’s centroid was defined as the mean across time of a trajectory in N-dimensional space with *N* = 365 for dSTR and *N* = 479 for lPFC.

### 2.2 Corticostriatal model

#### 2.2.1 Architecture

We jointly trained a connected system of two recurrent neural networks to perform the movement-sequence task (see Methods and Materials). Single-unit dynamics in these networks are governed by:

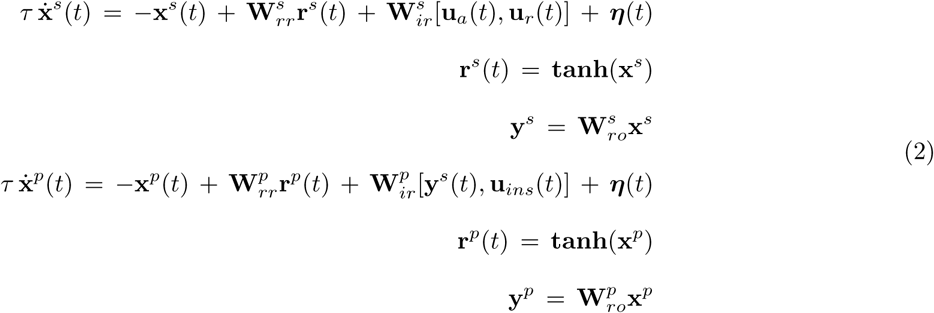

where 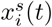 and 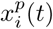 are synaptic current variables of unit *i* at time *t* in the striatal and prefrontal network, respectively, and activity (firing rate variables 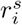 and 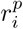) is a nonlinear function of x (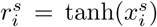 and 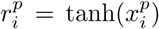), 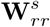 and 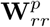 are the recurrent weight matrices, 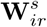 and 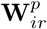 are the input weight matrices, and [**u**_*a*_, **u**_*r*_] and [**r**^*s*^, **u**_*ins*_] are the inputs of the striatal and prefrontal networks, respectively, and added noise *η*_*i*_(*t*).

The striatal and the prefrontal network had 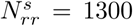 and 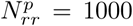 units, respectively. The connectivity weight matrices 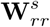 and 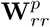 were initially drawn from the standard normal distribution and multiplied by a scaling factor of 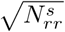 and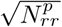, respectively. The neural time constant was *τ* = 10 *ms*. Each unit received an independent white noise input, *η*_*i*_, with zero mean and *SD* = 0.01. Inputs were fed into the networks through the input weight matrices 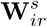 and 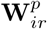 which were initially drawn from the standard normal distribution and multiplied by a scaling factor of 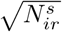 with 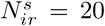 and 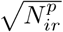 with 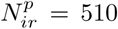 for the prefrontal network. Otherwise, the model parameters were set in the same range as [Mante et al., 2013].

The striatal network received a 15-dimensional input vector, **u**_*s*_ = [**u**_*a*_, **u**_*r*_], composed of a 10-dimensional vector **u**_*a*_ specifying actions taken and a 5-dimensional vector **u**_*r*_ specifying the rewards received for those actions. Outputs for each of the networks were linearly read out from the synaptic currents of the recurrent circuit (Eq 2). The striatal network read-out was a 10-dimensional vector, **y**^*s*^, of action values derived from TD learning (as described further below).

The prefrontal network received a 20-dimensional input, **u**_*p*_ = [**y**^*s*^, **u**_*ins*_], composed of a 10-dimensional vector **r**^*s*^ of action value outputs from the striatal network, together with a 10-dimensional instruction vector **u**_*ins*_ specifying when to initiate fixation (Fixate) and when to move towards or hold the target (Move + Hold). The prefrontal network read-out was a 11-dimensional output vector, **y**_*p*_ = [**y**_*a*_, *y*_*v*_] composed of a 10-dimensional vector **y**_*a*_ of actions and an additional unit coding for the visual hemifield (upper or lower) in which a particular sequence terminated.

#### 2.2.2 Network system training

All synaptic weight matrices (**W**_*rr*_, **W**_*ir*_, and **W**_*ro*_) were updated with the gradient of the loss function (Eq 3), which was designed to minimize the square of the difference between network and target output:

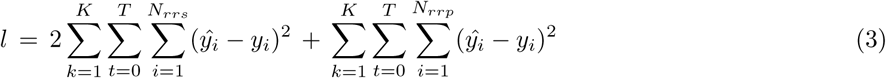

The error was thus obtained by taking the difference between network output *y* and target output *ŷ* and summing over all trials *K* in a batch (with *K* = 10), time points T and recurrent units *N*_*rr*_. The total loss function (Eq 3) was obtained by combining the loss terms from the striatal and the prefrontal network while assigning double weight to the striatal loss term. The striatal and the prefrontal networks were jointly trained by obtaining the gradient of the combined loss function (Eq 3) through automatic differentiation with *autograd* [Maclaurin et al., 2017] and custom implementations of specific functions with GPU-based acceleration using *JAX* [Johnson et al., 2018]. The network was trained with an initial learning rate of *α* = 0.001 for 10 steps with 1000 iterations each, while the learning rate was decayed by 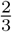 at every step. After this training phase, all the synaptic weight matrices remained fixed.

#### 2.2.3 Task Coding

Actions (**u**_*a*_/**y**_*p*_) and action values (**y**_*s*_) were coded as 10-D vectors in which every unit coded for one of the movement choice options (see Methods and Materials -Task and Stimuli). So, for instance, for the first binary choice option (Fig 1C, S1-center) one unit coded the right movement choice and the other the left movement choice. Rewards (**u**_*r*_) and visual inputs which indicated target locations and fixation points (**u**_*ins*_) were coded as 5-D vectors with every unit coding for reward delivered at one of the 5 decision points (center, upwards, downwards, upper, lower; Fig 1C). Actions and rewards were coded as brief transients. The reward signal interval was 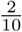 th as long as the action signal interval. Rewards were delivered after the end of the action signal interval.

Rewards drove the update of action values according to a temporal-difference reinforcement learning algorithm (Q-Learning) [Sutton and Barto, 2018]:

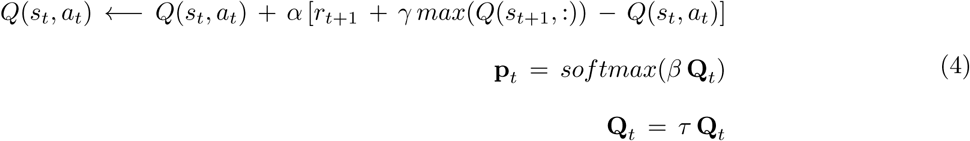

The learning rate parameter *α*, the discount factor *γ* and an additional inverse temperature parameter *β* were fitted to one of the training sessions of monkey 1 using *fminsearch* in Matlab, with the decay parameter set to *τ* = 0.95. The values obtained for the parameters were *α* = 0.8100, *γ* = 0.2010, and *β* = 3.050.

The training data set for the corticostriatal model system was composed of the behavioral data from the recording sessions of the two monkeys. Action values (**y**_*s*_) were derived by feeding the actions chosen and rewards received through the Q-learning algorithm (Eq 4). A subset of 25 blocks from the recordings was left out as a test set. Additionally, the training data set was augmented with artificial data generated by randomly choosing one of the eight sequences for the current block and drawing action movement outcomes with the same error probabilities that the animals displayed in the real task (see Fig. 4E -Behavior). During training, a batch composed of 10 blocks was randomly chosen from across the entire training set at each step.

**Fig 2.**
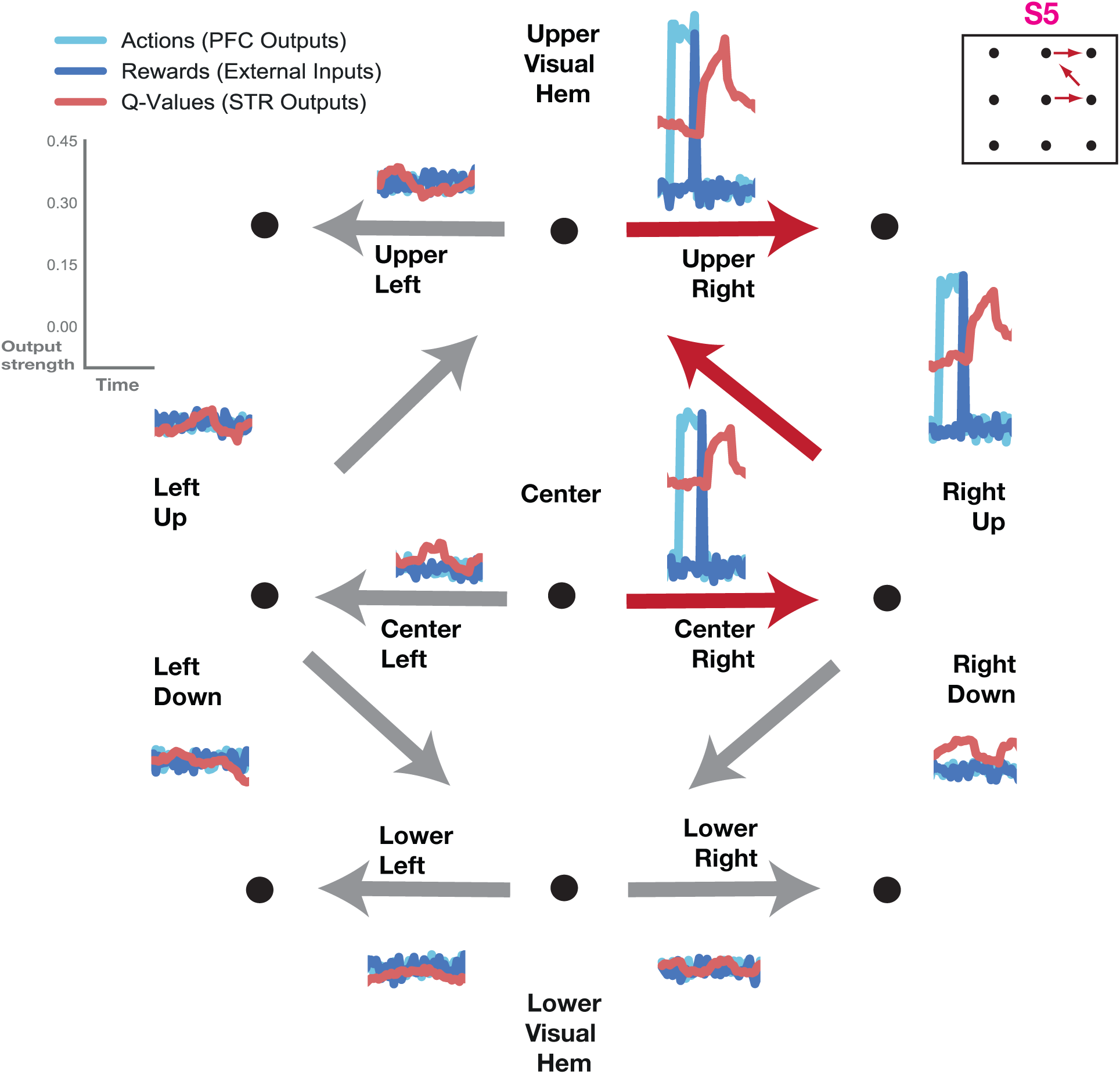
Task coding. The correct movement sequence (S5) was signalled by the corresponding output units while units coding other movement directions stay flat. Actions (light blue) were indicated by pulses in the prefrontal network’s output units corresponding to a particular direction. Rewards (dark blue, plotted on top) were delivered at the end of action pulses. Action values (red, plotted on top) were indicated by striatal output units corresponding to a particular direction; action values increased after reward delivery.

**Fig 3.**
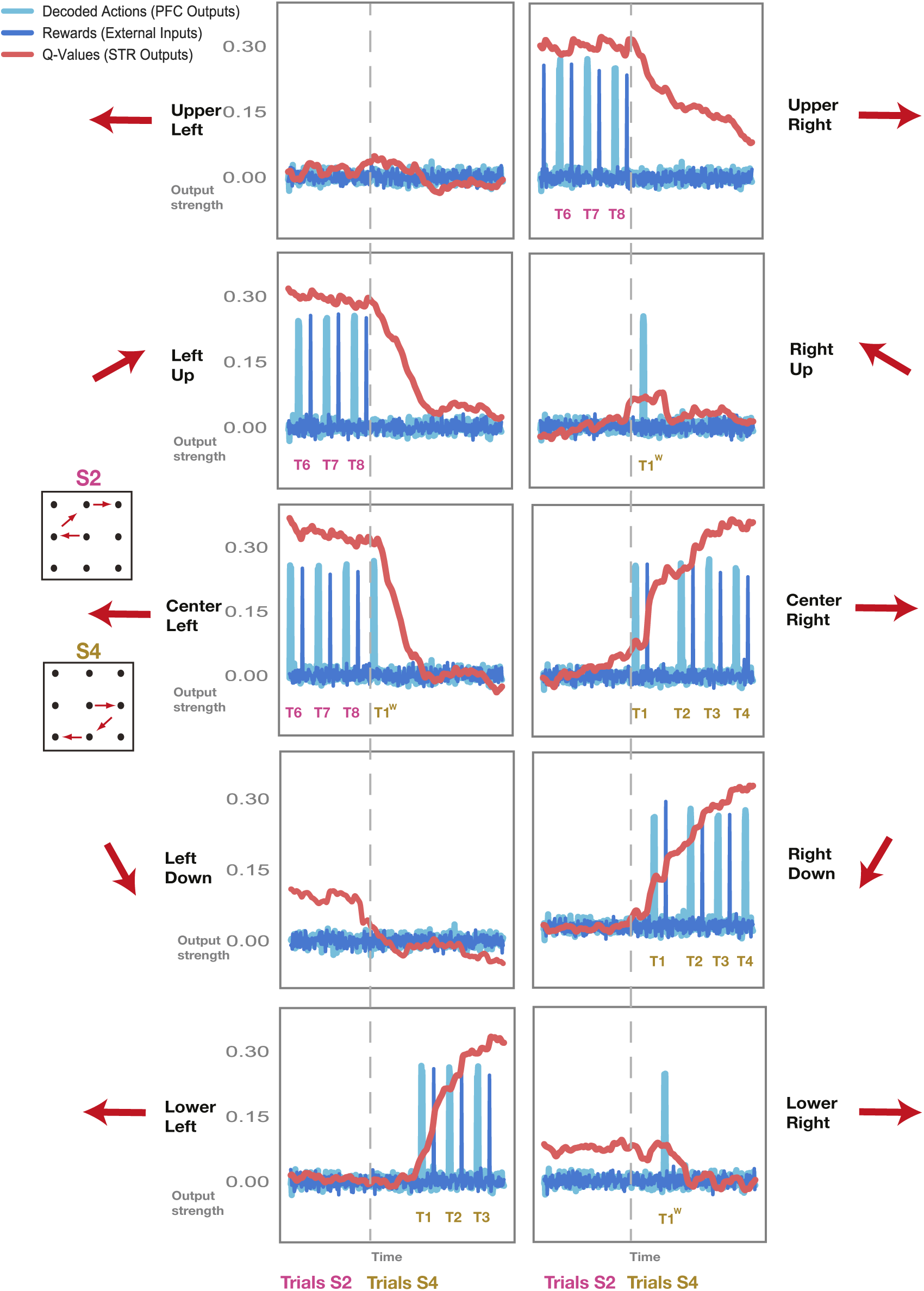
Corticostriatal model: autonomous behavior. Sample transition between two blocks in which the correct sequence switches (from S2, in magenta, to S4, in light brown; block transition indicated by vertical grey dotted line). Decoded Actions (light blue) are plotted together with rewards (dark blue) and action values (red) for all output units. Action values spike after correctly executed, rewarded actions and increase with successive correct actions. Action values for wrong, unrewarded moves decay quickly. Trials with erroneous movements remain unrewarded and are repeated, just like in the original experiment. Supporting Figures: Fig. S1

**Fig 4.**
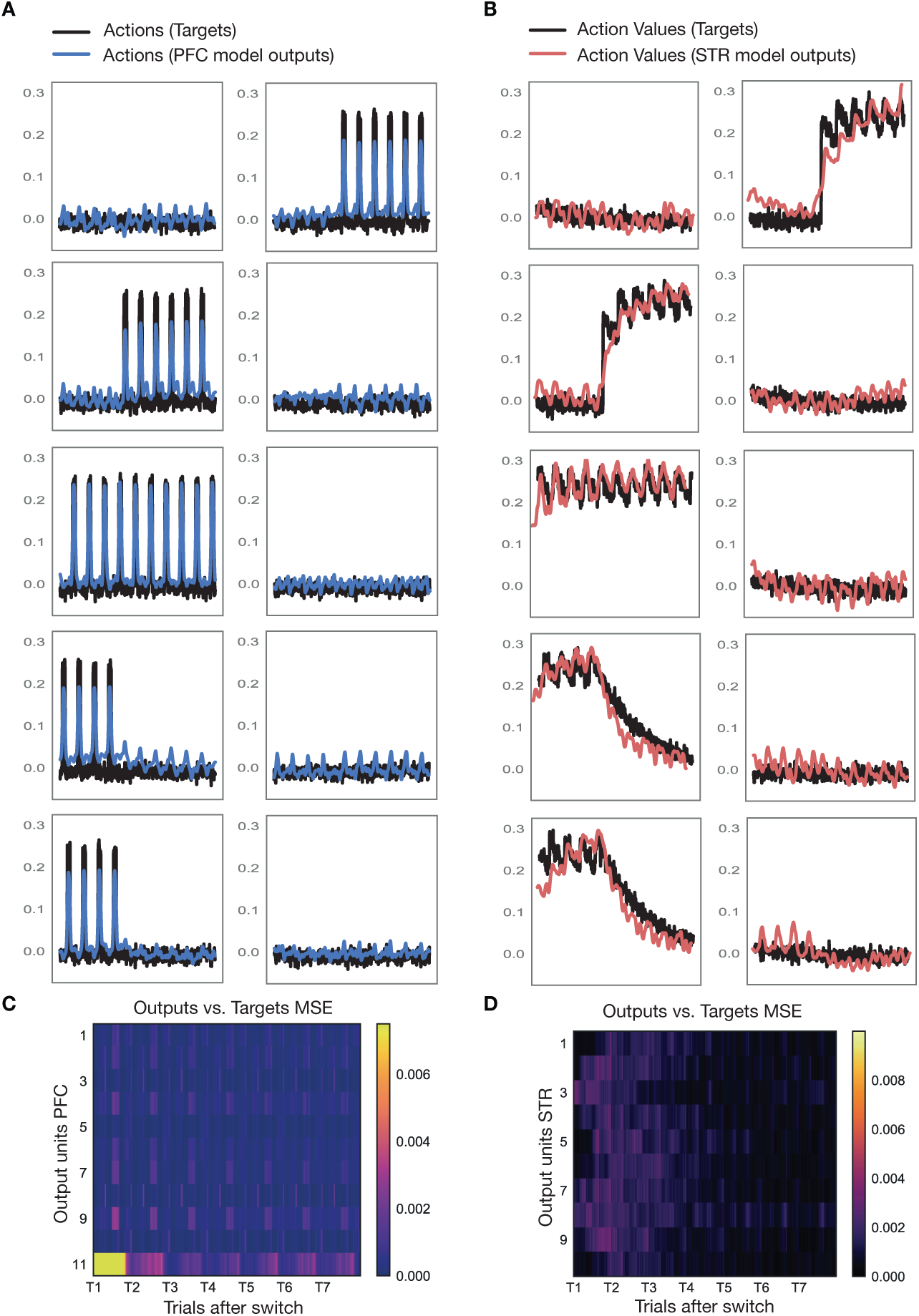
Model Performance. **A** Outputs of prefrontal model network (blue) versus targets (black) for a sample test block. **B** Outputs of striatal model network (red) versus targets (black) for the same sample test block as in A. **C-D** Mean squared error (MSE) between outputs and targets averaged over 25 test blocks. MSE is depicted as a function of trials after the sequence switched for all output units in the prefrontal (C) and striatal network (D). Supporting Figures: Fig. S2

#### 2.2.4 Autonomous mode

After training, we also tested the model’s ability to autonomously produce movement-sequence blocks. This meant obtaining trial-by-trial value and action estimates. More specifically, we obtained an initial set of outputs (**y**^*s*^; Eq. 2) from the striatal network by feeding in a vector of white noise inputs for **u**_*s*_ (with zero mean and *SD* = 0.01). Striatal outputs were then fed into the prefrontal network together with instructions **u**_*ins*_ specifying the particular movement stage as part of the prefrontal input vector **u**_*p*_. We obtained a vector of actions **y**_*p*_ as outputs from the prefrontal network. Actions were decoded from among the choice options that were possible at a particular movement stage (after the first movement stage, certain choice options become unavailable, i.e. for sequences S1 and S5 in Fig. 1C the choice options in the lower visual hemifield are no longer possible at the second movement stage). Actions were decoded probabilitiscally from the result of passing action outputs through a softmax function (**a**_*choice*_ = *softmax*(**y**_*p*_)).

The chosen action was then fed back into the dSTR network. If the action was correct, a reinforcement signal was delivered to the dSTR. If the action was incorrect, no reinforcement signal was delivered. Then, the signal for the next movement stage was fed into the dPFC network and a new set of dSTR outputs, for the next movement stage, was fed into the prefrontal network, etc.

If the decoded action was wrong (in that it was not the correct action for the current movement-sequence block), the network system was forced to repeat this particular movement stage before proceeding (just like the animals in the original experiment). This was done by concatenating the set of wrong actions with the striatal action input vector **u**_*a*_ and feeding it into the striatal network together with the reward vector **u**_*r*_ set to white noise. The striatal outputs obtained as such were then fed into the prefrontal network together with the instruction vector **u**_*ins*_ set to the same movement stage as before.

#### 2.2.5 Model Analysis

The neural population activity from the striatal and prefrontal networks of the corticostriatal model was imaged in the same way as the real neural recordings, by projecting activity into a 3-dimensional latent space spanned by the first three vectors of the *sequence*-decoding matrix (**D**_*s*_) obtained through dPCA (Eq. 1) [Brendel et al., 2011, Kobak et al., 2016].

Canonical correlation analysis (CCA) was used to compare model and neural population responses, as was done previously [**?**]. First, both the model and the recorded neural data were averaged across trials, smoothed with a Gaussian kernel and reduced to 15 dimensions using dPCA. This ensured CCA was not performed along dimensions of high correlation but low variance. Then the first 10 canonical correlation coefficients were obtained using the entire duration of a chosen movement-sequence as a comparison window.

In order to image the potential surface, we first projected neural population activity into 3-dimensional latent space and obtained a mesh of points (**X**^*^) around the sample trajectories. Then we projected this mesh of points back out into neural population space using the *stimulus*-encoder matrix (**F**_*s*_; Eq. 1). We started the networks off at each of these points by providing them as the initial vector of firing rates (**r(t)**; Eq. 2), and let the network system run through a whole block of movement-sequence trials. For illustration purposes, we imaged the potential surface at the same point for all trials in a block. We obtained the potential surface at the chosen moment in time by calculating the magnitude of the gradient for the mesh of points. In order to ensure that the trough of the potential surface at the chosen moments in time really pointed to fixed point regions, we kept the input fixed and continued running the network system through 10 iterations (which corresponded to the length of the fixation period) to make sure the location of the trough remained fixed on the timescale of the network. If the location of the minimum remained fixed for the length of this period, we labelled this location as a fixed point. We then obtained the joint potential surface of two chosen sequences by taking the point-wise minimum across the two sequences’ manifolds.

In order to calculate the distance between the minima of the potential surfaces of two different sequences, we first confirmed the location of the fixed points in 3-dimensional latent space as described above. Then, we used Dijkstra’s algorithm to obtain the minimal path length along the joint potential surface between two particular sequences’ fixed points. We repeated this for all possible sequence combinations in the test set and averaged the result. In order to calculate the distance between sequences in latent space as well as the distance of a particular sequence to its centroid, we first projected the data into 10-dimensional dPCA-space (for computational reasons) and then proceeded the same way as in the neural data (see Methods and Materials -Neural Data -Data Analysis). We found no difference in these metrics when projecting into 10-or 20-dimensional dPCA-space.

## 3 Results

We investigated dynamics during learning in the fronto-striatal system (Fig. 1). The task consisted of a sequence of three movements which the animal had to execute by saccading to the correct targets (Fig. 1A). While the animal exectued these movement trajectories, recordings were obtained from dPCA and lPFC (Fig. 1B). There were a total of 8 possible arrangements (or sequences) of 3 movements (Fig. 1C). These sequences, in turn, were arranged into blocks in which one particular sequence remained fixed for the duration of the block (Fig. 1D). Thus the animal was able to learn which sequence was correct in the current block using feedback about chosen movements. At the start of a new block, a new sequence was chosen randomly (see Methods and Materials for further details).

Two animals performed this task while we obtained neural recordings from lateral prefrontal cortex (lPFC) and dorsal striatum (dSTR) (Fig. 1B, see Methods and Materials and [Seo et al., 2012] for further details). To investigate learning dynamics, we built a model of the fronto-striatal system. In this model, consistent with the neural data, the striatum represents action values, and the prefrontal cortex selects actions (Fig. 1E). Whether actions are selected via return loops through the thalamus, or in other structures downstream from the striatum is currently unclear. However, at least in some conditions in the in-vivo study, actions were selected in cortex before they were selected in the striatum [Seo et al., 2012]. Therefore, we organized our model consistent with this. The prefrontal network received action value information from the striatum and selected actions based on this information. The prefrontal network received additional inputs signifying fixation and move/hold periods, which were presented to the animal as visual cues during the task. The two networks in this system were trained using actions and rewards from the animals’ real recorded behavior. Corresponding action values were generated by feeding actions and rewards through a temporal-difference reinforcement learning algorithm (Q-learning; see Methods and Materials).

After training, the fronto-striatal model learned to produce correct movement sequences (Fig. 2). In this example, sequence S5 (consisting of a rightwards movement, followed by upwards, and another rightwards movement) was the correct sequence for the current block. The prefrontal network units coding for these movements selected the correct action (light blue). Following selection of the correct action the network received a reward input (dark blue). The reward, in turn, made the value signal in the striatal network (red) increase for the rewarded movement. The output of the prefrontal and striatal network units coding for other movement directions remained flat. Altogether, the system learned to produce movement sequences with the correct action and action value output.

After training, the network system could be run autonomously. That is dlPFC selected actions based on the dSTR action value inputs, and external inputs to dlPFC indicating the ordinal position of the current movement in the sequence (see Methods and Materials -Corticostriatal Model -Autonomous mode). In the first step, a vector of white noise inputs and a vector of external inputs specifying the ordinal position of the movement were used to generate value signals in the striatal network. These value signals were input to the prefrontal network which selected the first movement. This movement was then fed back into the striatal network together with the corresponding reward outcome. The signal representing the new ordinal position in the sequence was used to generate the next value signal, and so on. In this manner, we generated several trials of the same movement sequence, which were strung together into blocks, as in the original experiment (Fig. 1D). We imaged the outputs of the network system as the correct sequence switched from one block to the next (Fig. 3). There were a total of 10 output units, one for each of the available movement directions. Decoded actions (in light blue) from the outputs of the prefrontal network, action value outputs (in red) from the striatal network and rewards (in dark blue) delivered externally were all imaged together for these two sequential blocks. The correct sequence switched from one block to the next (sequence S2, magenta, for the first block, and S4, light brown, for the second; Fig. 3 inlays).

The correct sequence for the first block (S2, magenta) is composed of a left move in the center, followed by an upwards move on the left side and a rightwards move at the top (see Fig. 3 inlay). Accordingly, the three output units at center left, left up and upper right were active. The correct actions could be decoded probabilistically from the outputs of the prefrontal network; actions were signalled by sequential pulses (light blue) in the three output units that made up the correct sequence. Rewards were presented as short pulses at the end of an action (dark blue). The striatal network subsequently produced the corresponding action value signal (red). The value signal decayed over time, but recovered when correctly executed trials followed upon each other (as in the upper right movement panel, Fig. 3).

As the new block began (grey dotted line), the correct sequence changed (from S2 in magenta to S4 in light brown). The new sequence was composed of a center right move, followed by a downwards move on the right and a move to the left at the bottom (see Fig. 3 inlay), ending up in the lower visual hemifield. Accordingly, the three output units at center right, right down and lower left were now active. The output units representing the movements that had been correct during the previous sequence (center left, left up, upper right) were now inactive; the striatal network’s value signal in these units decayed back towards zero.

Occasionally, the prefrontal network commits errors and the wrong movement was decoded. Errors occur with the same frequency as in the behavioral data from the two monkeys carrying out this task (as the training set for the networks was derived from real behavior, see Methods and Materials). Upon a block switch, errors can occur at all three movement stages. At the first movement stage, the PFC network produced the same movement which had been correct in the previous block (center left, T1^*w*^ in light brown; Fig. 3). This move remained unrewarded. The striatal value signal decayed quickly after unrewarded moves, instead of spiking as it did after rewarded moves (i.e. center left, T8 in magenta). The network was forced to repeat the move before progressing further (just like the animals were in the original experiment; see Methods and Materials). The striatal value signal for the opposite movement direction now increased, prompting the prefrontal network to produce the correct move at the second attempt (T1 in light brown, center right). The correct action was now rewarded and the striatal value signal for this move increased further, while the value signal for the unrewarded action decayed further (center left). Further erroneous moves occurred at the second and third movement stages (right up and lower right, respectively). These outputs remained unrewarded, and the network was again forced to repeat these moves. The correct moves were generated at the second attempt, and the network was allowed to progress to the next trial.

We also plotted the behavior of the network system on the test set (Fig. S1). In this case, actions from the test set (a portion of the original dataset which was not used in network training) were fed into the striatal network to obtain value signals which, in turn, were fed into the prefrontal network to obtain action outputs. Actions were not fed back into the striatal network this time; rather, the next action from the test set was picked as the next input to the striatal network. The network behaved similarly to the autonomous mode, with value signals increasing after successive correctly executed moves and decaying quickly upon the omission of reward (Fig. S1). To analyze the system’s performance in more detail, we plotted striatal and prefrontal outputs together with their target outputs for a sample block transition from the test set (Fig. 4A,B; see Methods and Materials). Output traces generally follow target traces. To quantify this, we computed the mean squared error between targets and outputs over the duration of a block averaged over all blocks in the test set (Fig. 4C-D). The largest difference between outputs and targets occurred for the first trial when the correct sequence was unknown. Over the course of the block, as certainty increased, the error decreased for both prefrontal (Fig. 4C) and striatal networks (Fig. 4D). We also imaged the eigenvalue spectra of the two networks’ weight matrices after training (Fig. S2). We found that variance was spread across many dimensions in the prefrontal network (Fig. S2A), while most of the variance was concentrated around 5 large eigenvalues in the striatal network (Fig. S2B).

We determined the behavior of the model system by measuring the fraction of correct decisions over the course of the movement-sequence block and compared it to the recorded behavior from the animals (Fig. 5A). The recorded behavioral data from the animals (solid line, as in [Seo et al., 2012]) shows chance performance at the beginning of the block, and rapid, steady improvement over the course of the block. In order to determine the fraction of correct decisions of the model system, we let the networks produce movement-sequences autonomously (see Methods and Materials-Corticostriatal model). That is, we used the action outputs of the prefrontal network as inputs to the striatal network. In this way we obtained a distribution of activations for the next movement step in the prefrontal output units, from which we decoded the predicted action. The fraction of correct decisions of the autonomous model system averaged over 25 blocks (dotted line) approximated that of the animals’ behavior (solid line). We also plotted the fraction of correct decisions obtained from the fit of the Q-learning algorithm (dashed line), and it agreed well with behavior. This shows the model system was able to capture the animals’ behavior in this task.

**Fig 5.**
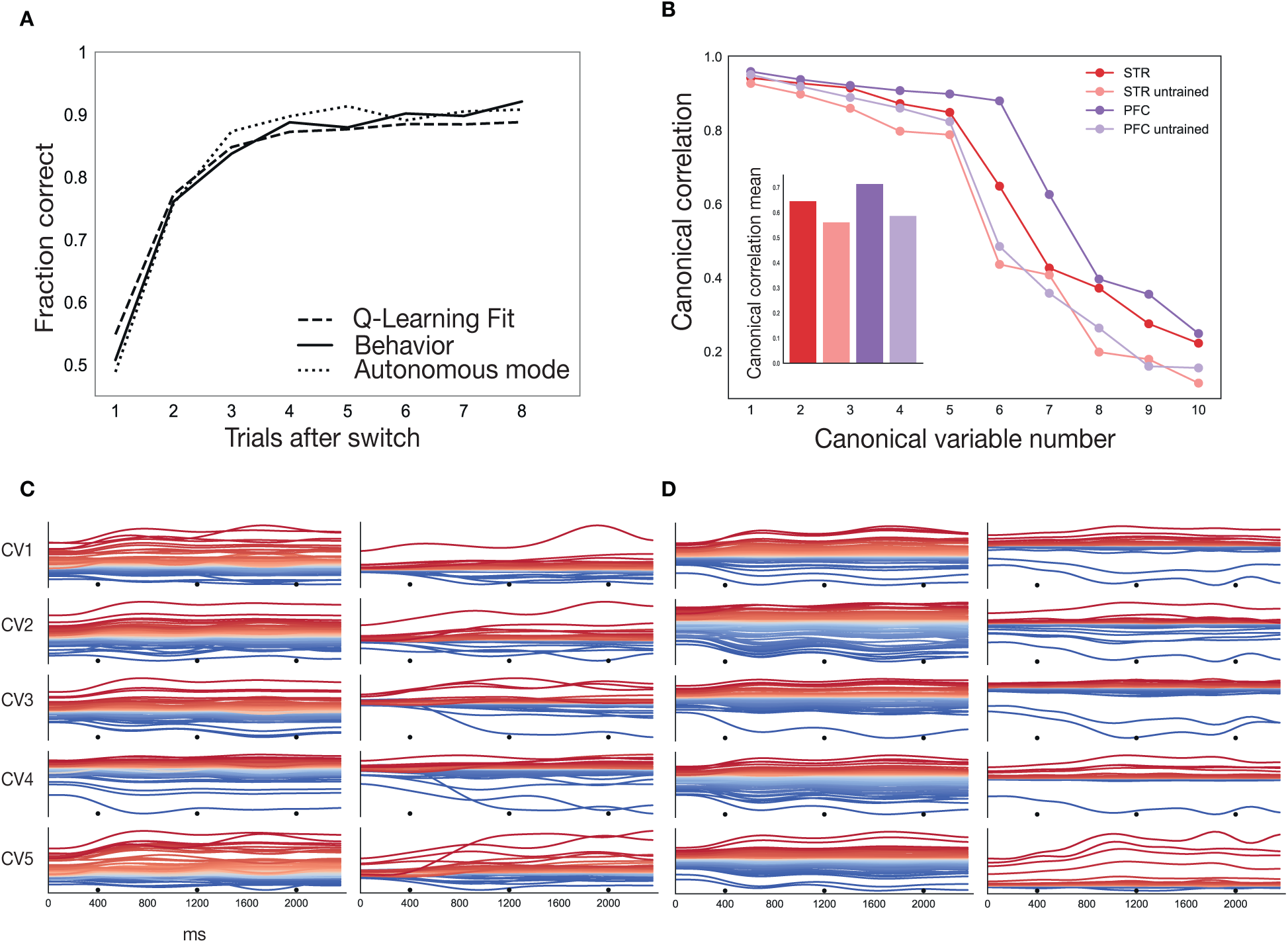
Comparison of model and recorded data. **A** Fraction of correct decisions as a function of trials after the sequence switched, separately for the behavior of the two animals (solid line), the performance of the Q-learning algorithm (broken line; see Methods and Materials-Corticostriatal model-Coding), and the performance of the corticostriatal model network system (dotted line; see Methods and Materials-Corticostriatal model-Autonomous Mode). **B** Summary of canonical correlations between model and neural data. CCA analysis provides a spectrum of of correlation coefficients that can be used to assess model performance (see Methods and Materials). The canonical correlation coefficients are shown for the trained model (striatal network in dark red and prefrontal network in dark magenta), as well as for an untrained network with the same inputs which as a baseline (striatal network in light red, and prefrontal network in light magenta). **C** CCA projections (canonical variables) for the *striatal* model network (left) and the neural data recorded from dSTR (right). These projections show the directions in state-space along which the data is maximally correlated with the model. Each row shows one canonical variable (CV 1-5). Traces are colored based on the mean value of the projection across the entire depicted duration of the trace. Traces show responses for the whole length one movement-sequence which is composed of three sequential movements (black dots mark movement onset). **D** CCA projections for the *prefrontal* model network (left) and the neural data recorded from PFC (right).

We also compared neural activity from the model with the recorded neural data using canonical correlation analysis (Fig. 5B; see Methods and Materials). We plotted the canoncial correlation coefficient for the striatal and the prefrontal network across the first ten canonical variables. We also obtained the correlation coefficients for an untrained model with the same inputs as a baseline. The canonical correlations were consistently higher than the baseline for both the striatal and the prefrontal network. The first seven correlation coefficients were significant for the trained striatal and prefrontal networks (*p* < 4.823*e* − 11). The average correlation coefficients were higher for the trained networks (0.65 striatal and 0.71 prefrontal) than for the untrained networks (0.56 striatal and 0.59 prefrontal). We also plotted the first five canonical variables for the neural responses from the model and the recorded data (Fig. 5C,D). The canonical variables are the directions in state-space along which model and recorded data are maximally correlated, and offer a way to assess the similarity between model and recorded population responses. The model (left) and the neural data (right) shared many population-level response patterns. The model was able to pick up the slow oscillatory patterns observed in the neural data [Seo et al., 2012].

To study how neural representations evolved with learning, we imaged PFC neural population activity in 3-dimensional latent space using demixed principal component analysis (see Methods and Materials). Trajectories from neural recordings in lPFC (Fig. 6A-D) are plotted alongside trajectories from the prefrontal network of the corticostriatal model (Fig. 6E-H). The plots capture representations for progressive trials over the course of a block (Fig. 1D), as the animals advance from 50% certainty at the start of the block (Fig. 6A&B) to close to 90% certainty by the fifth trial into the block (Fig. 6C&G).

**Fig 6.**
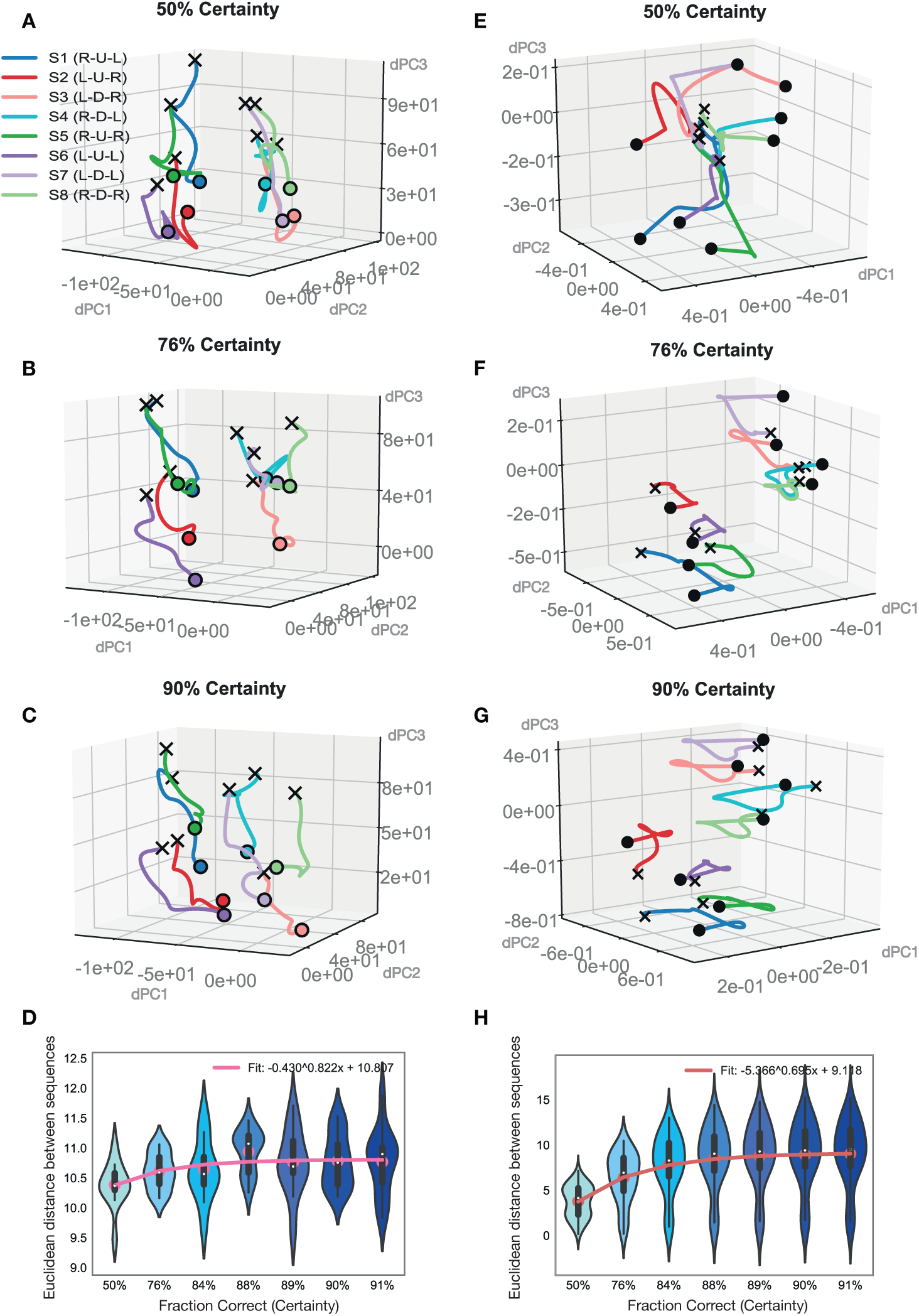
Evolution of latent task coding with learning for PFC recordings and model network. **A-D** lPFC neural recordings. Latent task trajectories during the execution of the three-movement sequence depicted in three-dimensional latent space (obtained from dPCA using the sequence dimension; see Methods and Materials-Neural Data-Data Analysis). Latent trajectories for all eight possible movement sequences (S1-S8) are plotted together, for increasing certainty (fraction correct) levels (A-C). **D** Euclidean distance between all sequence trajectories within the two clusters (brightly and lightly colored trajectories) as a function of increasing certainty (see Methods and Materials-Neural Data-Data Analysis). **E-H** PFC model network. Latent task trajectories during the execution of the three-movement sequence depicted in three-dimensional latent space (obtained from dPCA using the sequence dimension, as in A-D). Latent trajectories for all eight possible movement sequences (S1-S8) are plotted together, for increasing certainty levels (E-G). **H** Euclidean distance between all sequence trajectories within the two clusters (brightly and lightly colored trajectories) as a function of increasing certainty level (calculated as in D).

Sequence representations from neural recordings in lPFC (Fig. 6A-D) showed a separation by visual hemifield: sequences S1, S2, S5 and S6 which progressed along the upper visual hemifield (Fig. 1C) were clustered to the left while the remaining sequences which progressed along the lower visual hemifield were clustered to the right. This separation appeared to be present from the very start of a block. We also examined the evolution of representations across learning. As the animals learned in each block, they more frequently selected the correct option (Fig. 4E), which suggests they become more certain of their choices. We found that sequence trajectories within each particular cluster separated more from each other with increasing certainty (Fig. 6A-C). To capture this effect, we computed the Euclidean distance between sequences within clusters in neural population space (in the full-dimensional space of recordings, not in the reduced latent space; see Methods and Materials). We found this measure increased with certainty as learning progressed over the block (Fig. 6D). Sequence representations from the prefrontal network model (Fig. 6E-H) showed the same clustering by visual hemifield. Task trajectories were somewhat more distinctly clustered by hemifield at the start in the neural data (Fig. 6A) than in the model (Fig. 6E). Overall, though, we also found Euclidian distance between sequences within clusters increased for the model trajectories (Fig. 6H).

To study what underlies increasing separation of movement-sequence representations with learning, we probed the prefrontal model network further (Fig. 7). We obtained the potential surface in the region around the latent sequence trajectories as learning progressed across the block (see Methods and Materials -Model Analysis). We obtained the joint surface across two particular movement-sequence trajectories in latent space by taking the point-wise minimum across the potential surfaces of the individual sequences, and imaged this common surface for increasing levels of certainty across a block (Fig. 7A-C). The potential surface shows how activity evolves at different points in neural latent space in the vicinity of sequence trajectories. Neural activity has a high propensity to be pushed away from locations where the magnitude of the gradient is high (yellow), and remain in locations where the magnitude is low (dark blue). We ascertained that the troughs of the potential surface pointed to energy minima (or fixed points) in neural activity (see Methods and Materials -Model Analysis).

**Fig 7.**
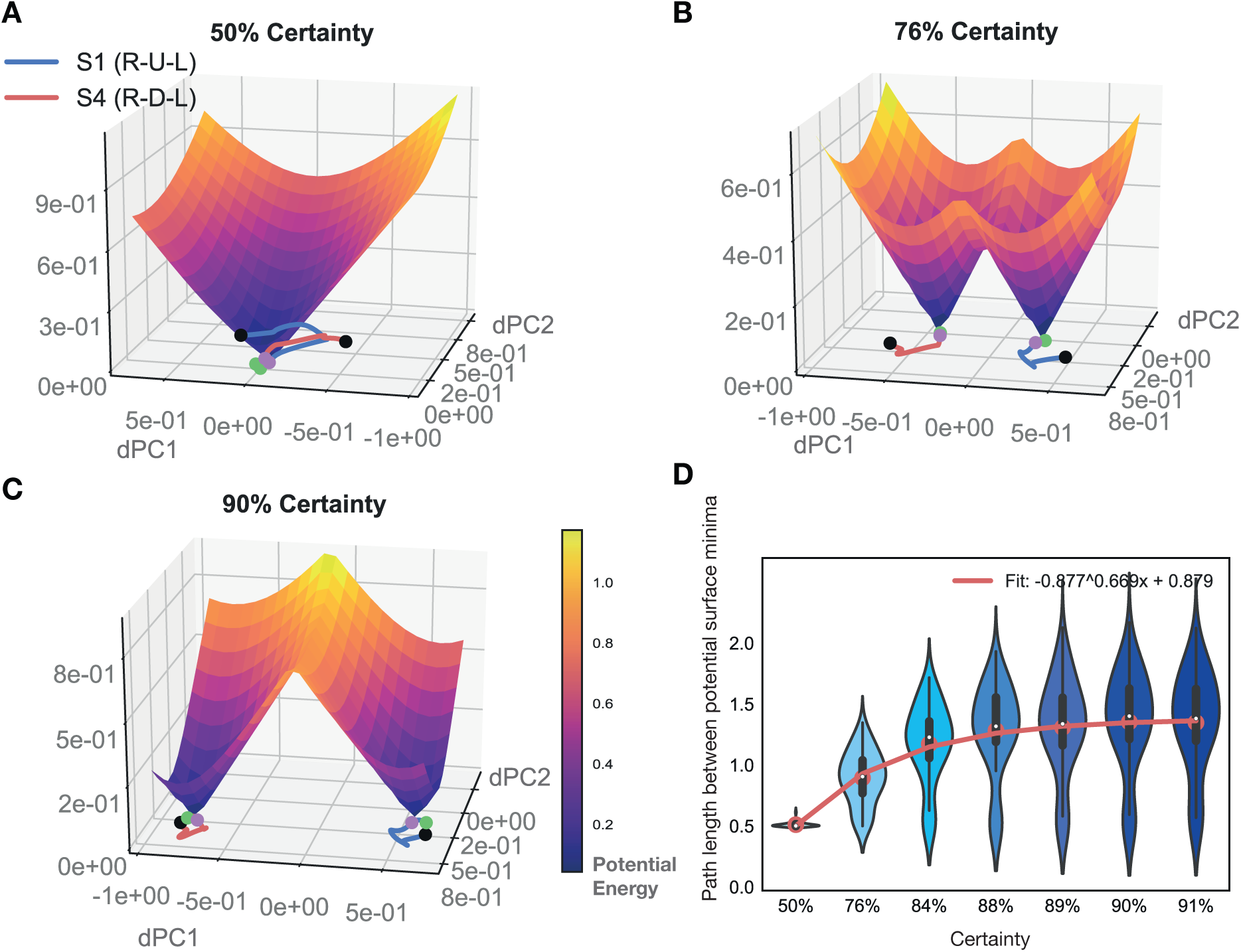
Evolution of potential surface with learning in PFC model network. **A-C** Latent trajectories (obtained through dPCA) for two different movement sequences are depicted in two dimensional latent space (S1 in blue and S4 in red, on the bottom). Solid dots depict the beginning (green) and end (black) of the three-movement sequence. The surface depicts the potential energy of the network in the two dimensional space in which the two sequence trajectories sit. To obtain the potential surface, the network was iterated one step forward with inputs held fixed to a particular chosen timepoint (magenta dot; see Methods and Materials-Corticostriatal Model-Model Analysis). Latent movement-sequence trajectories and potential surface are depicted for increasing certainty (fraction correct) levels in A-C. **D** Minimum path length between the gradient minima of all sequence pairs in the test set. Path length was calculated along the joint gradient surface by using Dijkstra’s algorithm (see Methods and Materials-Corticostriatal Model-Model Analysis). Results are depicted for increasing certainty levels.

Observing the evolution of joint potential surfaces with learning (Fig. 7A-C) for a particular point along the movement trajectory (magenta dot), one notices how energy minima for different regions become increasingly well separated. Along with this separation, the ridge in the joint potential surface between the two sequence minima heightened with increasing certainty during learning. As this happens, it becomes increasingly less likely to commit errors: the gradient along a particular side of the ridge drives activity more strongly towards a particular sequence’s fixed point region, so that the chance to end up in the other minima decreases with learning. To quantify this effect, we determined the minimal path length between the various sequence’s fixed point regions as learning progressed (see Methods and Materials -Model Analysis). We observed that minimal path length between fixed point regions increased with certainty during learning (Fig. 7D). Altogether, we established that the energy landscape in the network changes with learning such that fixed point regions for different movement-sequences are pushed farther apart from each other, underlying the increase in behavioral accuracy.

We also examined how neural representations evolved with learning in the striatum (Fig. 8). Trajectories from neural recordings in dSTR (Fig. 8A-D) are plotted alongside trajectories from the striatal model network (Fig. 8E-H). Sequence representations in the dSTR (Fig. 8A-D) did not display any particular clustering by visual hemifield, unlike representations in lPFC (Fig. 6A-D). Sequence representations were instead scattered around in latent space. As learning progressed, sequence representations spread further apart from each other (Fig. 8A-C). We computed the Euclidean distance between all sequences in neural population space and found it to be increasing with learning (Fig. 8D). Trajectories from the striatal model network (Fig. 8E-H) were also scattered around latent space, similar the neural recordings (Fig. 8A-D). Like in the recordings, sequence representations in the model spread further apart from each other as learning progressed. To quantify this effect, we again computed the Euclidean distance between sequences (see Methods and Materials) and found it to be increasing with learning (Fig. 8H), as in the neural data (Fig. 8D).

**Fig 8.**
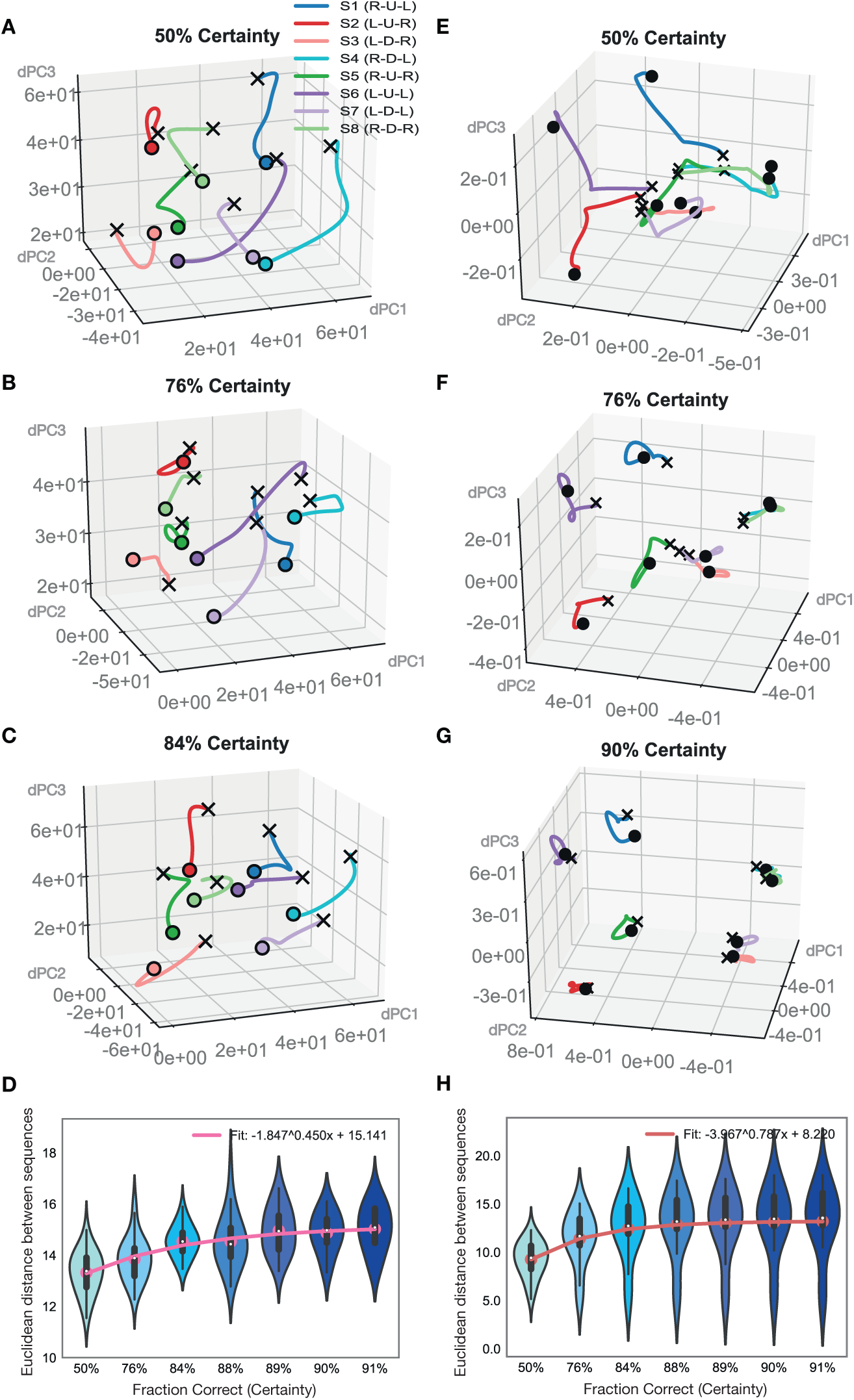
Evolution of latent task coding with learning for STR recordings and model network. **A-D** dSTR recordings. Latent task trajectories during the execution of the three-movement sequence depicted in three-dimensional latent space (obtained from dPCA using the sequence dimension; see Methods and Materials-Neural Data-Data Analysis). Latent trajectories for all eight possible movement sequences (S1-S8) are plotted all together for increasing certainty (fraction correct) levels (A-D). **D** Euclidean distance between all sequence trajectories within the two clusters (brightly and lightly colored trajectories) as a function of increasing certainty level (see Methods and Materials-Neural Data-Data Analysis). **E-H** STR model network. Latent trajectories for all eight possible movement sequences (S1-S8) are plotted together in dPCA-derived latent space (obtained as in A-D) for increasing certainty (fraction correct) levels (E-G). **H** Euclidean distance between all sequence trajectories within the two clusters (brightly and lightly colored trajectories) as a function of increasing certainty level.

To examine this effect in further detail, We computed the potential surface for various points as learning progressed (Fig. 9), as done previously for the PFC model. In the STR model network we also observed energy minima for different sequences becoming increasingly well separated as learning progressed (Fig. 9A-C). Along with this separation, the ridge in the joint potential surface between different sequence minima heightened with increasing certainty during learning. To quantify this effect, we determined the minimal path length between the various sequence’s fixed point regions as learning progressed, as before. We observed that minimal path length between fixed point regions increased with certainty during learning in the STR model network (Fig. 9D). Altogether, we established that the energy landscape in the network changes with learning such that fixed point regions for different movement-sequences are pushed farther apart from each other.

**Fig 9.**
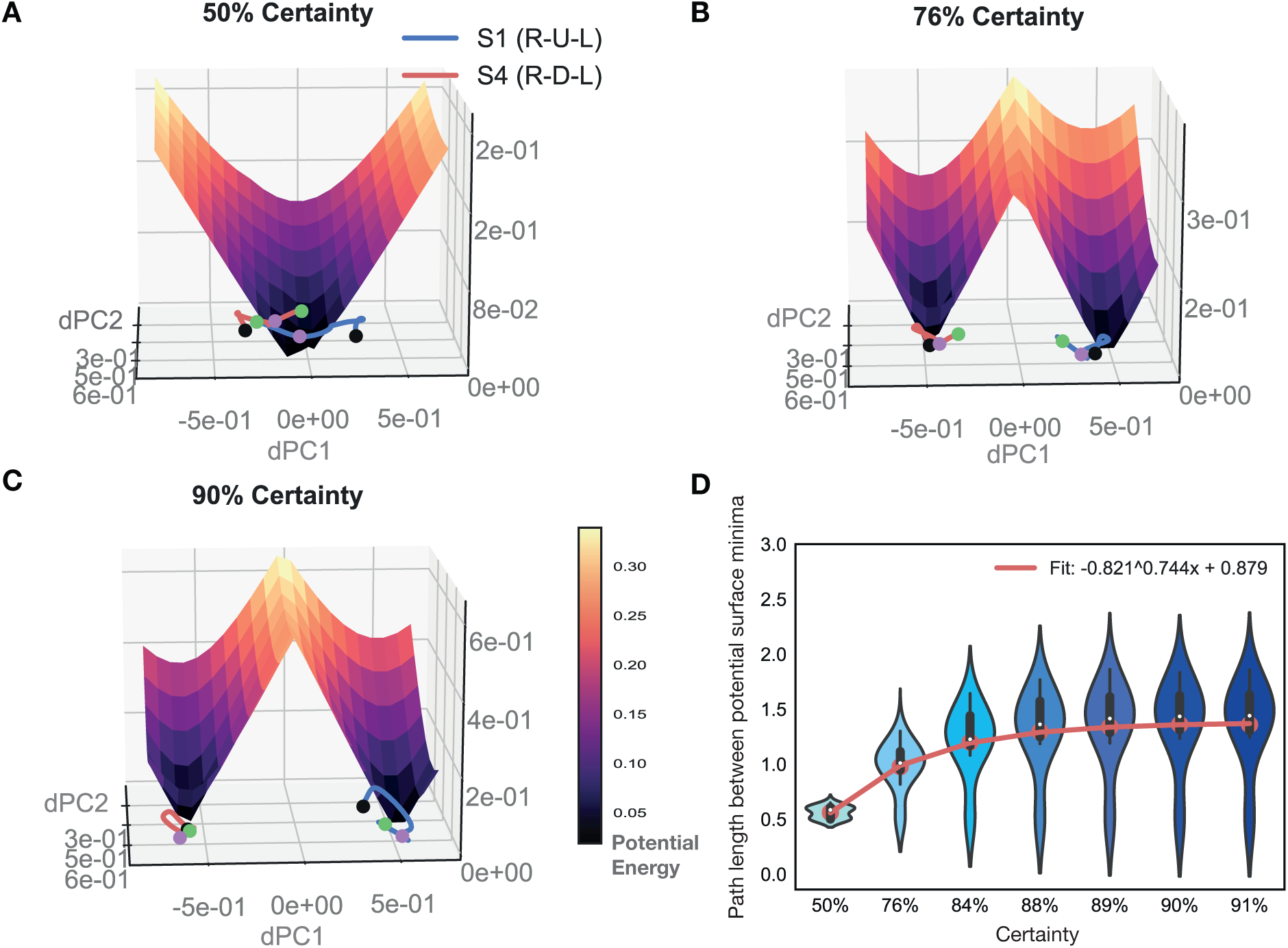
Evolution of gradient landscape with learning in STR model network. **A-C** Latent trajectories (obtained through dPCA) for two different movement sequences (S1 in blue, S4 in red) are depicted in two dimensional latent space (bottom). Solid dots depict the beginning (green) and end (black) of the three-movement sequence. The surface depicts the gradient of the network in the two dimensional space in which the two sequence trajectories sit. To obtain the gradient manifold, the network was iterated one step forward with inputs held fixed to a particular chosen timepoint (magenta dot; see Methods and Materials-Corticostriatal Model-Model Analysis). Latent movement-sequence trajectories and gradient surfaces are depicted for increasing certainty (fraction correct) levels (50%, 75% and 90% certainty in A-C). **D** Minimum path length between the gradient minima of all sequence pairs in the test set. Path length was calculated along the joint gradient surface by using Dijkstra’s algorithm (see Methods and Materials-Corticostriatal Model-Model Analysis). Results are depicted for increasing certainty levels.

We also examined how the shape of a particular movement-sequence representation in latent space changes with learning in the two regions (Fig. 10). We observed that trajectories became more compact with learning (from lighter to darker blue) as could be seen in neural recordings from dSTR (Fig. 10A) and in the striatal model network (Fig. 10B). To quantify this effect, we computed the Euclidean distance of a particular sequence to its centroid for increasing certainty levels (see Methods and Materials) and found this measure to be decreasing with learning in both striatal recordings (Fig. 10C) and in the striatal model (Fig. 10D). The decreasing trend in prefrontal recordings was not significant (Fig. 10E), while our PFC model network showed a more clear decreasing trend (Fig. 10F).

**Fig 10.**
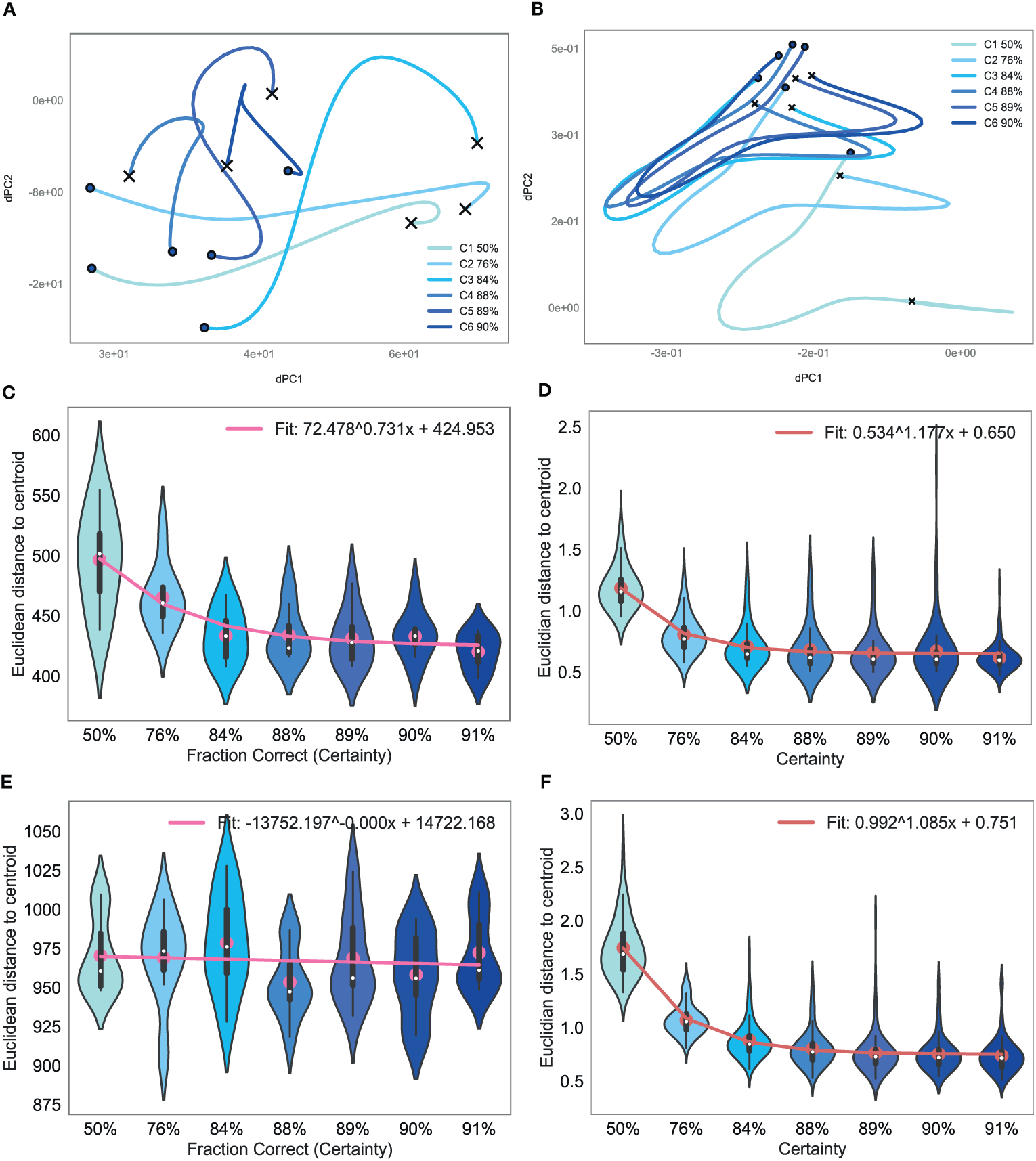
Shaping of latent movement sequence representations with learning. **A-B** Latent trajectory for a single movement sequence (obtained through dPCA) depicted in two dimensional latent space for increasing certainty levels (from low, in light blue, to high in dark blue). Neural recordings from dSTR (A) and STR model network (B). **C-F** Euclidean distance between a sequence’s centroid (multi-dimensional mean across time) and each point in time for all sequences in the test set (averaged across time points), as a function of increasing certainty (fraction correct). Neural recordings from dSTR (C) and STR model network (D). Neural recordings from lPFC (E) and PFC model networks (F).

We further compared latent representations for correct and wrong movements in the prefrontal model network (Fig. 11). We imaged representations of the first movement during the first trial in the block (when error rate is highest) in 3-dimensional dPCA-derived latent space for a few different sample sequences. We found that when the wrong movement was executed (i.e. S1 Error, dotted blue line), the representation moved away from that of the correct movement (right move, R in bold, for sequence S1, solid blue line) and closer to that of sequences which shared the same executed movement (left move, L in bold, for sequence S2, solid red line, and sequence S3, solid rose line). Similarly, the movement representation for the wrong move in sequence 8 (S8, dotted light green line) moved away from the correct move trajectory (solid light green line) and closer to the movement representation of sequence 6 (solid magenta line) which shared the same movement direction.

**Fig 11.**
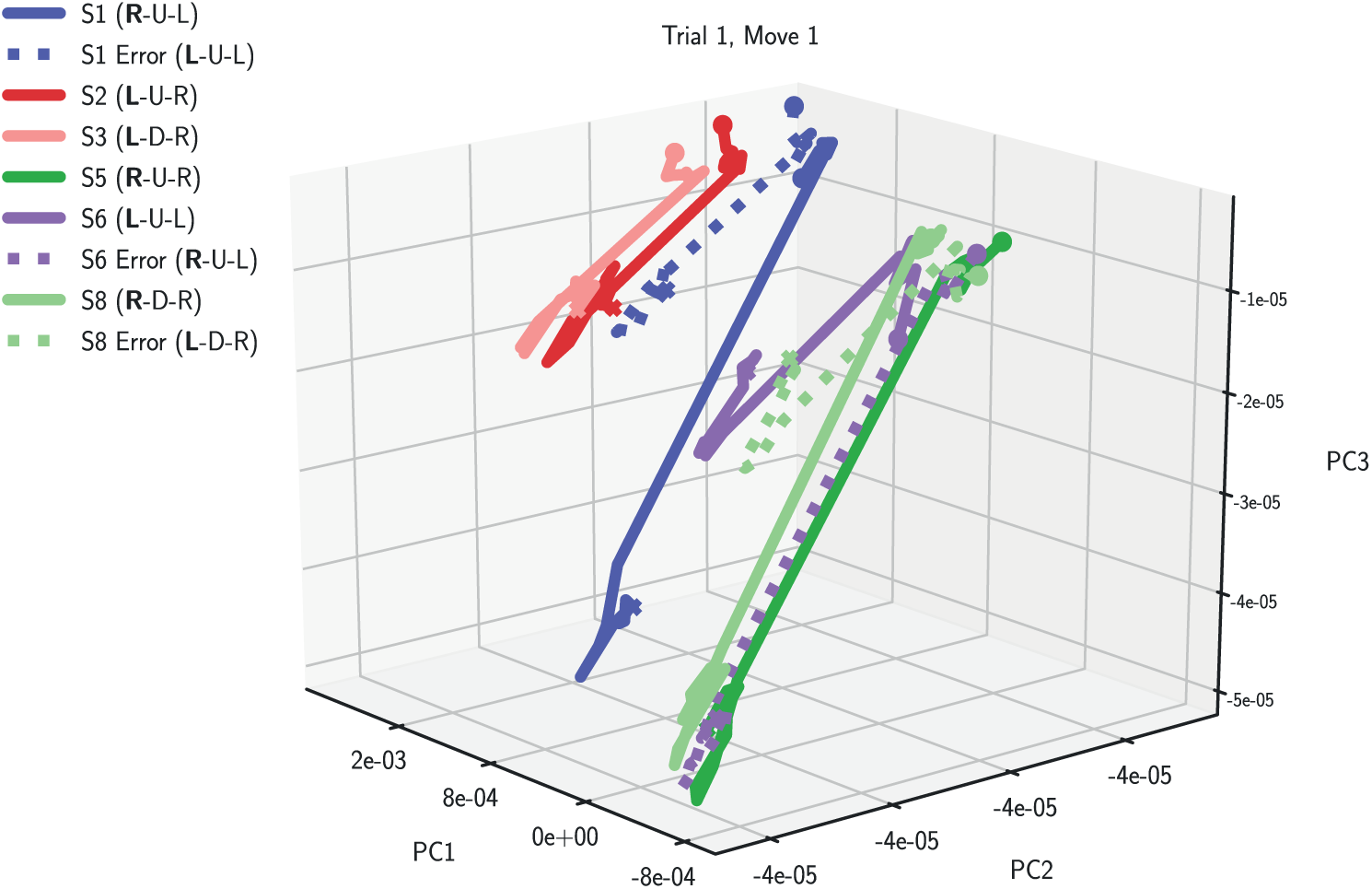
Latent representations of correct and wrong movements in the STR model network. Latent trajectories of single movements (obtained through dPCA) from the first trial after a sequence switch imaged in three dimensional latent space. Representations for correctly executed movement representations (solid lines) are depicted alongside wrongly executed ones (dotted lines).

## 4 Discussion

The results reveal how learning shapes neurophysiological responses in the corticostriatal system in the brain. Examining data from recordings in the dlPFC-dSTR circuit during an oculomotor sequence learning task, we found that learning increased the distance between action-sequence representations in activity space. To examine this further, we built a model of the corticostriatal system composed of a striatal network encoding action values and a prefrontal network selecting actions. This model was able to autonomously perform the task, matching the animals’ behavioral accuracy and closely approximating the representational structure present in neural recordings. Our model revealed that learning shapes the gradient landscape such that fixed point regions corresponding to different action sequences are pushed farther apart from each other. This makes it more likely the network generates the correct action. This offers testable predictions: when task learning is poor, representations should be more clustered together in activity space, making accurate decision making and decoding difficult. Altogether, this work shows how learning is expressed at the network level and suggests network level disruptions may lead to improper task learning.

Our model and results agree with previous findings from the corticostriatal system [Pasupathy and Miller, 2005, Samejima et al., 2005, Seo et al., 2012]. We found that task representations in neural recordings from lPFC were organized by visual hemifield, unlike dSTR representations, which showed no particular organizational structure. lPFC representations were also found to be less malleable with learning; the increase in distance between fixed point regions was smaller than in the dSTR. This agrees with previous findings that show less units in lPFC coding for a learning signal than in dSTR ([Seo et al., 2012]). Seo et al found less units in lPFC that coded for a reinforcement learning (RL) variable and more that encoded sequence information. Conversely, more units in dSTR showed a significant effect for RL and less units showed an effect for sequence information.

We also found that striatal representations became more compact with learning, both in recordings and in our model. This happened at the same time as sequence-specific representations spread further apart from each other in activity space. This effect could be partially driven by changes in the mean and variance of the neural population firing rate with learning, as well as by changes in higher order statistics. A decrease in responses among top-down signals as rewards become more predictable fits well within a predictive coding framework [Summerfield and de Lange, 2014, Keller and Mrsic-Flogel, 2018]. Previously it was found that the Fano factor, a measure of variability (variance of spike count divided by its mean), decreased in prefrontal neurons with learning [Qi and Constantinidis, 2012]. Also, changes in firing patterns within the neural population with learning -such as changes in synchronous firing [Baeg et al., 2007] -might be responsible for the changes in latent representation observed here.

Representations in recordings from lPFC were not found to become significantly more compact with learning, unlike in dSTR. This could be a consequence of less units in lPFC encoding a learning signal [Seo et al., 2012]. In our model we did not see a difference in how compact the representations became with learning in the two networks. This could be amended by additional processing steps which transform the output of the striatal network before it reaches the prefrontal network, which were not included in our model.

Our corticostriatal model system respects neuroscientific evidence that implicates the striatum in action value representation [Pasupathy and Miller, 2005, Samejima et al., 2005, Averbeck and Costa, 2017] and the PFC in action selection [Averbeck et al., 2006]. The methods used to train this system, however, are not biologically validated, similar to previous approaches [Mante et al., 2013, Yamins et al., 2014, Chaisangmongkon et al., 2017, Yang et al., 2018]. Alternatively, one could implement this system with spiking networks [Nicola and Clopath, 2017]. Another approach may be to use a reinforcement learning paradigm, rather than the gradient of an error signal, to train the network system [Song et al., 2017]. However, the latent dynamics underlying task learning uncovered here evolve over a longer timescale than the learning dynamics underlying network training, so findings are likely not impacted by different training protocols.

Our findings are in line with previous work showing that activity in a large population of neurons is confined to a lower dimensional manifold [Mante et al., 2013, Shenoy et al., 2013, Chaisangmongkon et al., 2017, Remington et al., 2018, Wang et al., 2018, Gallego et al., 2018]. We found striatal and prefrontal responses encoded the sequence task on a low-dimensional manifold. Previous work was confined to investigating dynamics on this manifold once the task was acquired (and the manifold fixed). We investigated how learning shaped the manifold during task acquisition, and found that learning is expressed in dynamics acting upon that manifold. These dynamics re-shaped the manifold in such a way that it became less likely for the network to commit errors.

## 5 Conclusion

We used a model of the corticostriatal system together with recordings from dlPFC-dSTR circuit in macaques during an oculomotor sequence learning task to investigate how learning shapes neural responses at the population level. The corticostriatal model was able to autonomously perform the task and to approximate neural task representations. Probing the model, we found that learning shapes latent representations such that it becomes less likely to commit errors as learning increases; the potential surface is reshaped with learning such that fixed point regions representing different action choices move farther apart from each other. All in all, this work shows how neural circuit dynamics in the corticostriatal system drive task learning.

## 6 Acknowledgments

This work was supported by a Wellcome Trust NIH-fellowship (C.D.M., S.S. % B.B.A.) and by NIH grant ZIA MH002928-01 (B.B.A.).

## 8. Supporting information

**Fig S1.**
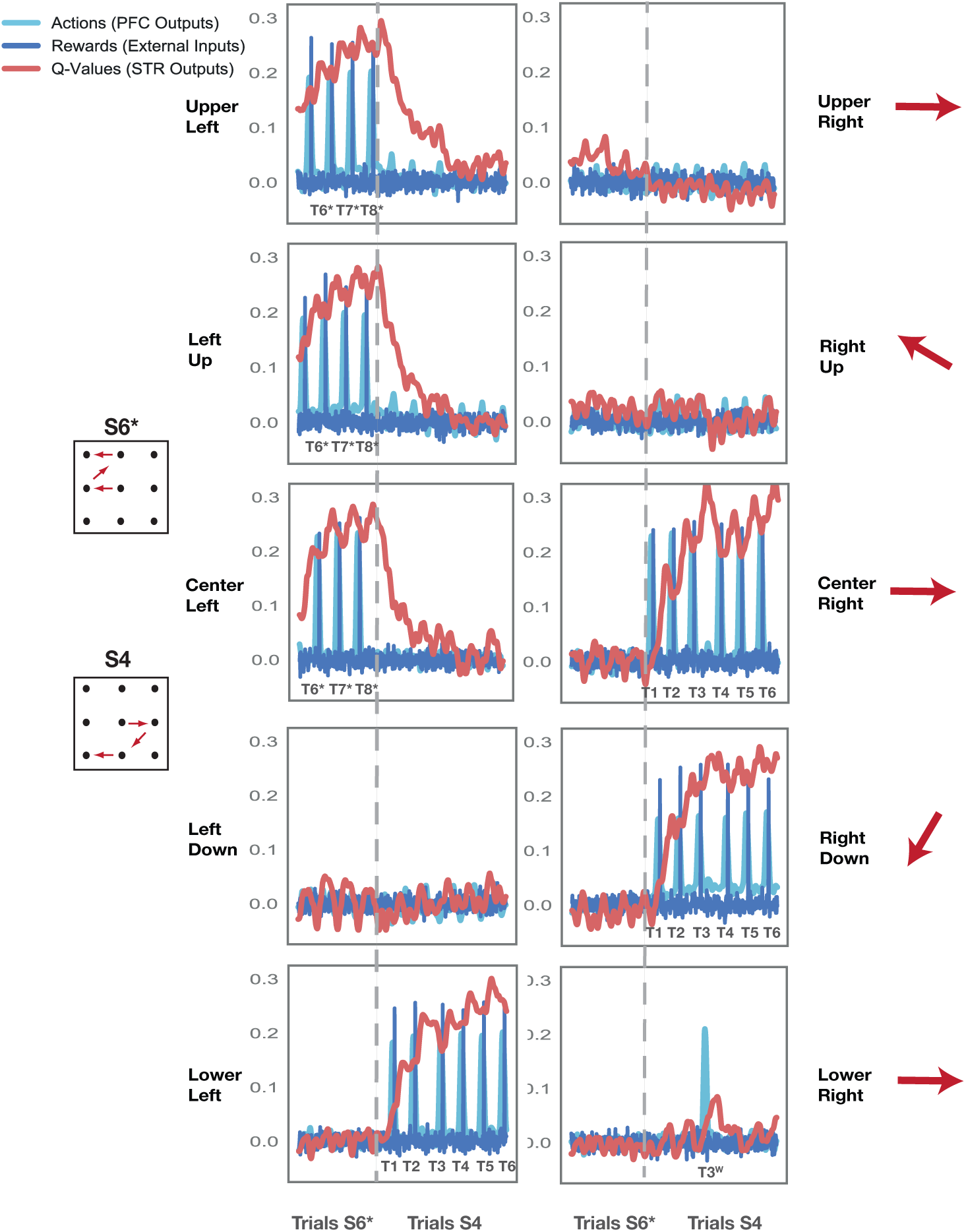
Corticostriatal model behavior on test set. Sample transition between two blocks in which the correct sequence switches (from S6 to S4; block transition indicated by vertical grey dotted line). Actions (light blue; outputs of the prefrontal (PFC) network) are plotted together with rewards (dark blue; external inputs) and action values (red; outputs of the striatal (STR) network) for all output units. Action values spike after correctly executed, rewarded actions and increase with successive correct actions. Action values for wrong, unrewarded actions (i.e. on the second trial of the third move, bottom right plot) increase briefly but decay quickly, never reaching the height associated with correct moves.

**Fig S2.**
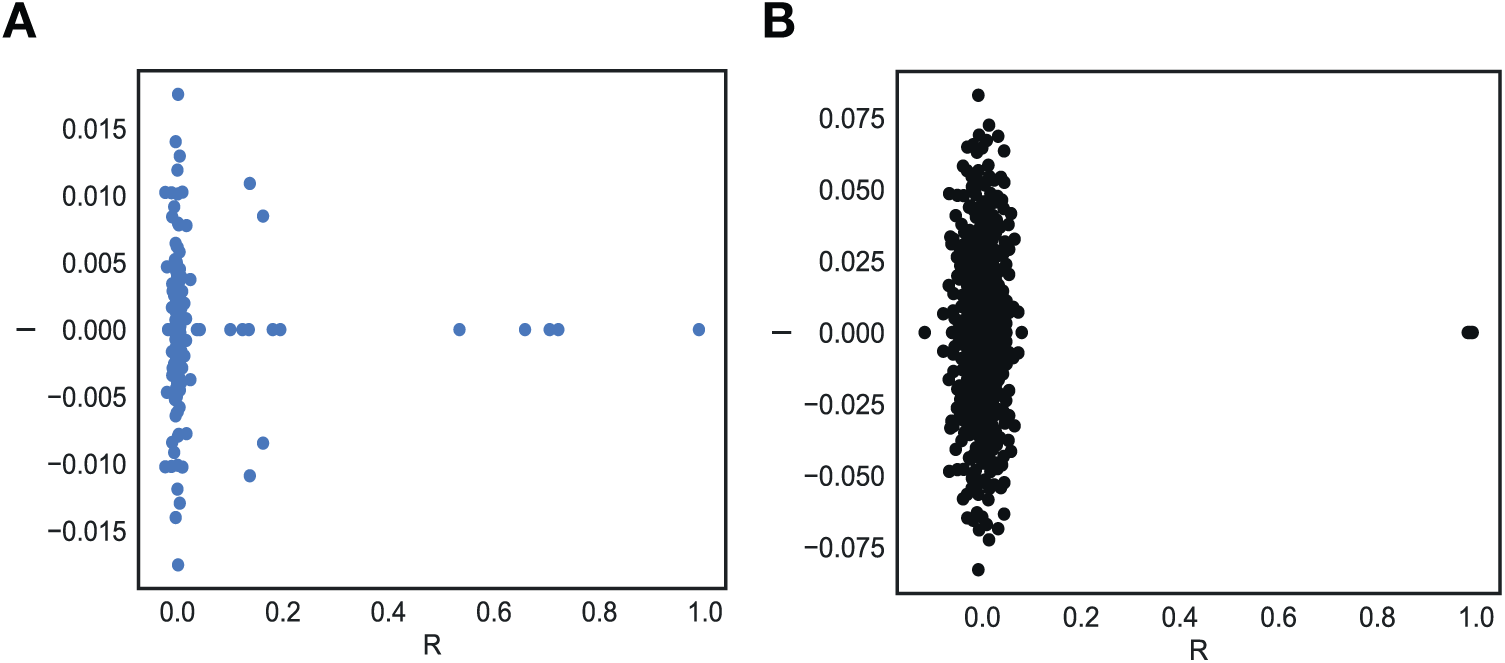
Eigenvalue spectra after training of the network system. **A** PFC model network, **B** STR model network.

